# Complete assembly of a dengue virus type 3 genome from a recent genotype III clade by metagenomic sequencing of serum

**DOI:** 10.1101/204503

**Authors:** Chitra Pattabiraman, Mary Dias, Shilpa Siddappa, Malali Gowda, Anita Shet, Derek Smith, Barbara Muehlemann, Krishnapriya Tamma, Tom Solomon, Terry Jones, Sudhir Krishna

**Author notes:** These authors contributed equally to the work (Chitra Pattabiraman, Mary Dias). Corresponding author: Chitra Pattabiraman.

## Abstract

**Background:** Mosquito-borne flaviviruses causing diseases such as dengue and Japanese encephalitis are devastating, particularly in the tropics. Although, multiple flaviviruses are known to co-circulate in India, when a patient presents with febrile illness, testing is usually limited to specific pathogens. Unbiased metagenomic sequencing of febrile cases can reveal the presence of multiple pathogens and provide complete genome information. Sequence information, a cornerstone for tracing virus evolution, is relevant for the design of vaccines and therapeutics. In order to assess the usefulness of unbiased metagenomic sequencing for the identification of viruses associated with febrile illness, we sequenced serum from four individuals and plasma from one individual, all hospitalized at a tertiary care centre in South India with severe or prolonged febrile illnesses, together with one healthy control in 2014.

**Results:** We identified and assembled a complete dengue virus type 3 (DENV3) sequence from the serum of a case classified as severe dengue. We also found a small number of Japanese encephalitis virus (JEV) sequences in the serum of two adults with febrile illness, including the one who had dengue. Phylogenetic analysis of the dengue sequence indicates that it belongs to a predominantly Asian, DENV3, genotype III clade. It had an estimated divergence time of 13.86 years (95% Highest Posterior Densities 12.94 - 14.83 years) with the closest Indian strain. Amino acid substitutions were present throughout the sequenced genome, including 11 substitutions in the antigenic envelope protein compared to the strain used for the development of the first commercial dengue vaccine. Of these one substitution (E361D) was unique and six were in critical antigenic sites.

**Conclusions:** We demonstrate that both genome assembly and detection of a low number of viral sequences are possible by unbiased sequencing of clinical material. Complete dengue virus sequence analysis places the sequenced genome in a recent, predominantly Asian clade within genotype III of DENV3. The detection of JEV, an agent not routinely tested in febrile illness in India, warrants further analysis and highlights the need to study co-circulating flaviviruses in parallel.

## BACKGROUND

Acute undifferentiated febrile illness refers to a sudden onset of high fever without localized organ specific clinical features [1]. Although majority of the cases recover over a few days, some can develop severe illness resulting in high morbidity and even death in many parts of the world. Among the many causes of febrile illness, some of the most important across Asia are mosquito-borne viruses such as dengue virus [1–6]. In addition, novel agents associated with acute febrile illness continue to be discovered [7–9].

Current molecular diagnostics, such as polymerase chain reaction are pathogen specific, and therefore pose limitations, as they may fail to detect co-infections and novel agents, not commonly associated with the disease syndrome [10]. Unbiased metagenomic sequencing of clinical material from patients with acute fever can overcome these limitations [3, 11].

Mosquito-borne viruses of the family *Flaviviridae* which include dengue and Japanese encephalitis viruses are known to co-circulate in India and other parts of Asia [12]. Dengue viruses are a major cause of acute febrile illness in Asia with recurrent outbreaks [13]. Japanese encephalitis virus on the other hand, is better known as a cause of acute encephalitis [14]. Although it has been noted as an agent causing acute fever in southeast Asia, it is not routinely tested as a cause of fevers in India [5, 6]. There are four distinct serotypes of dengue viruses (dengue virus 1-4), with small RNA genomes (approximately 10.8 kilo base pairs), making them amenable for characterization by deep sequencing of infected mosquitoes or clinical material from infected individuals [15]. Sequencing dengue genomes is important for tracking virus evolution, given that they frequently mutate [15, 16]. Outbreaks of severe dengue disease associated with serotype switches or introduction of a novel strain into the population have been reported from different countries, including Sri Lanka, Pakistan and Singapore [17–22]. Recent analysis suggests an Influenza virus like pattern for dengue virus evolution, where strain-specific differences underlie antibody neutralization [23].

Preexisting antibodies to circulating dengue strains can therefore contribute to disease severity by inadequate neutralization of the virus or by antibody mediated enhancement which facilitates virus infection [24–28]. This is supported by in vitro studies in which changes to the envelope protein of dengue virus type 3 (DENV3) were sufficient to alter antibody binding [26]. Multiple dengue vaccines are currently in various stages of development and a tetravalent vaccine (CYD-TDV; Dengvaxia^®^, Sanofi Pasteur) has been approved for use in several countries [29, 30]. This vaccine has been shown to induce broadly neutralizing antibodies to multiple strains and all serotypes of dengue viruses[31]. The results of the phase III trial of this vaccine suggest that both the immune state (with respect to dengue viruses) and circulating viruses may influence vaccine effectiveness [29]. This underscores the need to characterize both the sequence evolution and antibody response of circulating dengue strains.

We hypothesized that the unbiased sequencing or metagenomic approach would help us determine both the identity and the sequences of viruses in febrile illness. In particular, based on previous studies of sequencing data from the serum of febrile individuals, we expected that medium depth sequencing (about 10-20 million sequence reads per sample) was necessary and sufficient for providing complete sequences of small viral genomes from clinical material [2, 9]. In order to test this, we sequenced serum from four individuals and plasma from one individual presenting with febrile illness at a tertiary care hospital in Bangalore, India and one healthy control, during the dengue season of 2014. We recovered the complete coding sequence of DENV3 clustering into a recent genotype III clade.

## RESULTS

We sequenced RNA extracted from the serum of four patients hospitalized with severe febrile illness, and one plasma sample from a patient hospitalized with prolonged febrile illness (Table 1). We included serum from a healthy individual, and water as controls. Approximately 10×10^6 sequence reads were recovered from each sample, with the water control yielding lower number of reads (Figure 1 A).

**Table 1.**
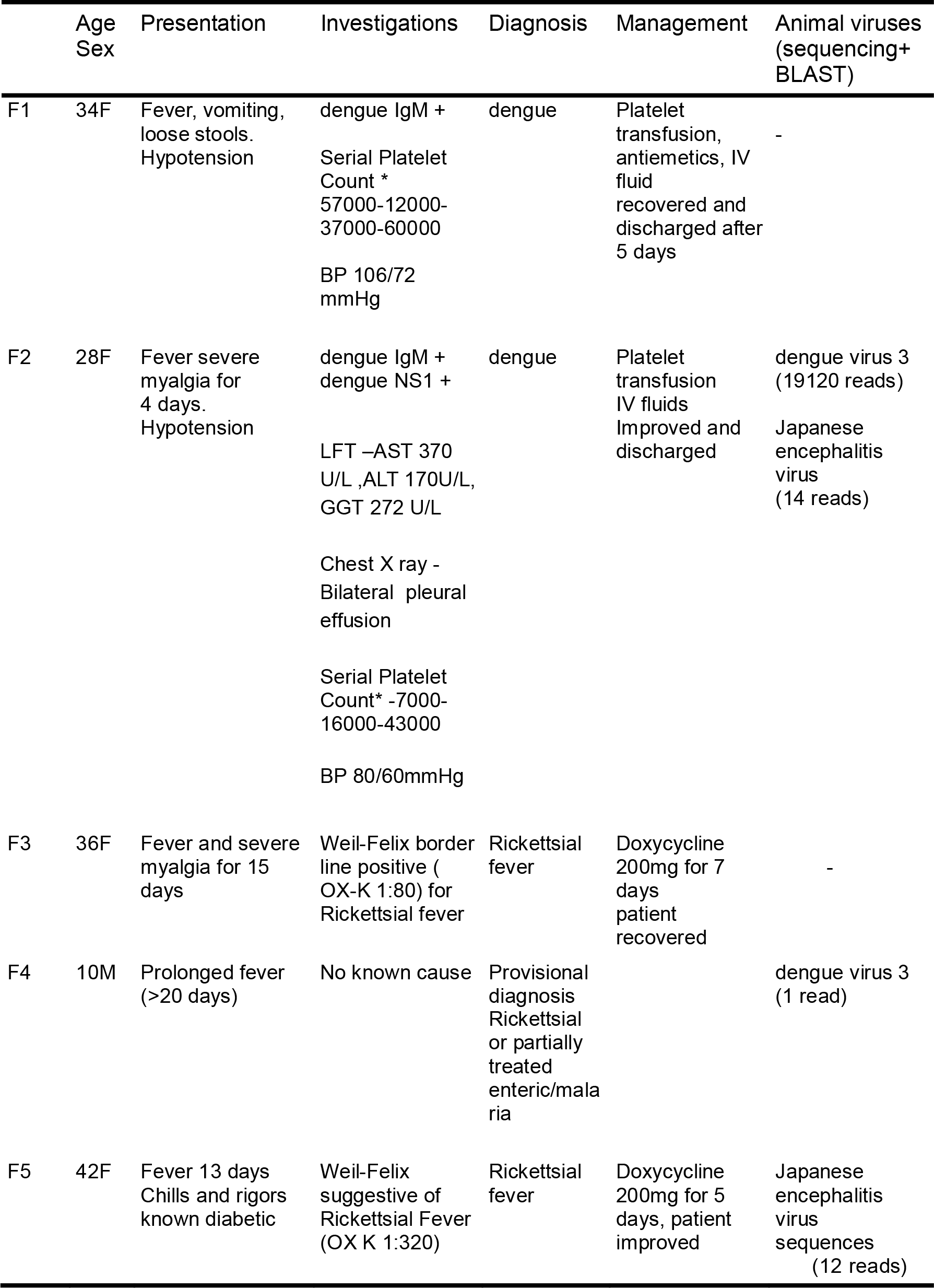
Clinical Profile of the sequenced cases. – Table shows the clinical presentation, key diagnostics tests, provisional diagnosis, treatment followed and results from sequencing (SNAP alignment against viral databases). Abbreviations used - M - Male, F -Female, IgM - dengue Immunoglobin M, NS1 - dengue Non-Structural protein 1 test, LFT- Liver function test, AST- Aspartate amino transferase, ALT- Alanine amino transferase, GGT- Gama Glutamyl transferase, *cells/mm^3^. “-” in column 7 indicates that no sequences mapping to viruses of animals (Animal Viruses) were confirmed by nucleotide BLAST in that sample. Numbers in brackets represent the number of sequence read that aligned to that virus using the SNAP alignment tool.

**Figure 1:**
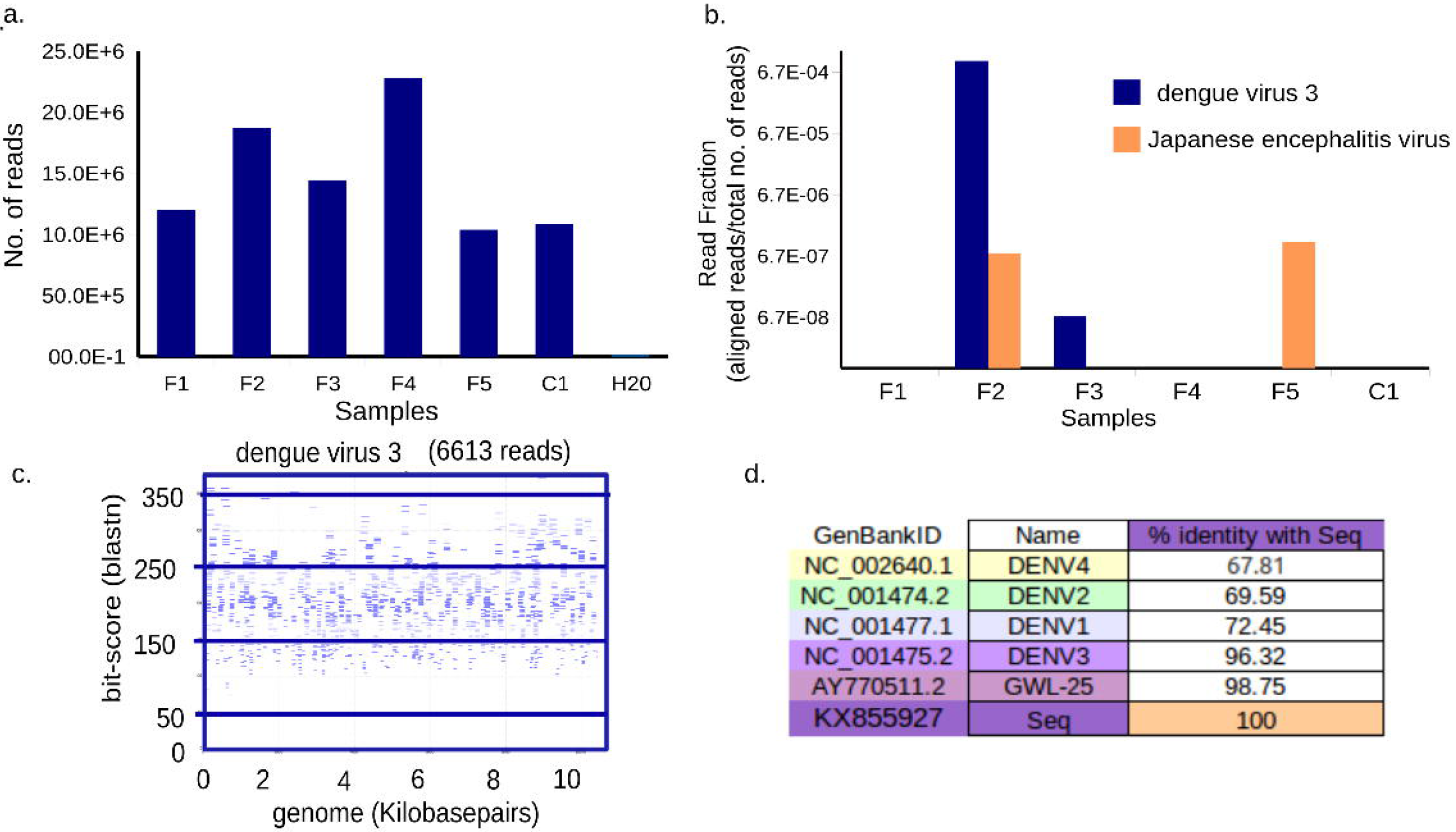
Dengue virus type 3 (DENV3) and Japanese encephalitis virus sequences identified from febrile serum 1a. Shows number of sequence reads generated per sample. 1b. Bar graph shows number of reads that aligned to a particular virus as a fraction of the total number of reads (Y axis, log scale) from that sample (X axis) using the SNAP alignment. 1c. Shows the alignment of sequences mapping only to DENV3 by nucleotide BLAST. Each rectangle shows sequencing reads (blue lines), their alignment to the genome (X axis) and their blast bit-score (Y axis). Numbers below the title represents number of reads that mapped to the title. 1d. Shows the percentage identity of KX855927 with all 4 dengue viruses and the closest Indian strain.

A BLAST [32] similarity search mapping of all sequenced reads to a database of NCBI reference viral sequences (Table 1) identified 19,120 DENV3 sequence reads and 14 Japanese encephalitis virus (JEV) sequence reads in sample F2, and 12 JEV sequence reads in sample F5. A single DENV3 read was detected in sample F3. No viruses of animals were confirmed by BLAST in the controls or in other samples (Table 1 and Figure 1B).

Based on the World Health Organization guidelines for classification of dengue cases [33], F2 was classified as a case of severe dengue as the presenting symptoms included respiratory distress (bilateral pleural effusions in chest X-ray) hypotension and elevated liver enzymes (Table 1).

The serum sample from this individual was positive for both the NS1 antigen and dengue IgM, and we were able to obtain complete DENV3 genome sequence from this sample. Genomes were assembled both by de novo assembly (87.05% coverage) and mapping based assembly (99% coverage) (Table 2-3, Additional File 1) and found to be identical (Additional File 2). Mapping revealed good coverage across the genome with an average depth of 231.45 (Figure 1, Table 2). The genome is missing 76 bp at the 5' UTR and 28 bp at the 3'UTR compared to the NCBI RefSeq (NC_001475.2) DENV3 genome.

**Table 2.** Assembly characteristics for mapping based assembly. - Table shows the quality, coverage and percentage nucleotide identity of the assembled DENV3 genome using different back bones and sequences for mapping using MIRA assembler. Backbone = Reference genome used for assembly, av.qual = Average quality of assembly, #-reads = Number of reads in assembly, mx.cov = maximum coverage of assembled genome by reads, av.cov= average coverage of assembled genome by reads, GC% = Percentage GC content of assembly, No cov = number of nucleotides of reference not covered in assembly.

| Criteria | Backbone | av.qual | #-reads | mx.cov. | av.cov | GC% | CnNoCov |
| --- | --- | --- | --- | --- | --- | --- | --- |
| All Reads from F2<br>against all 4 Refseq<br>of dengue viruses | DENV3 | 41 | 2009 | 96 | 26.27 | 46.67 | 145 |
|  | DENV1 | 30 | 2 | 3 | 1.01 | 46.67 | 10587 |
|  | DENV2 | 30 | 1 | 1 | 1 | 45.82 | 10723 |
|  | DENV4 | 30 | 1 | 1 | 1 | 47.12 | 10649 |
| "virus reads" from<br>F2 against all 4<br>Refseq of dengue<br>viruses | DENV3 | 42 | 18180 | 788 | 231.53 | 46.66 | 104 |
|  | DENV1 | 30 | 3 | 4 | 1.02 | 46.67 | 10587 |
|  | DENV2 | 30 | 1 | 1 | 1 | 45.82 | 10723 |
|  | DENV4 | 30 | 1 | 1 | 1 | 47.12 | 10649 |
| "virus reads" from<br>F2 against DENV3<br>and an Indian strain | DENV3 | 42 | 18178 | 793 | 231.51 | 46.66 | 104 |
|  | AY770511.2 | 43 | 18696 | 792 | 236.58 | 46.65 | 104 |

**Table 3.**
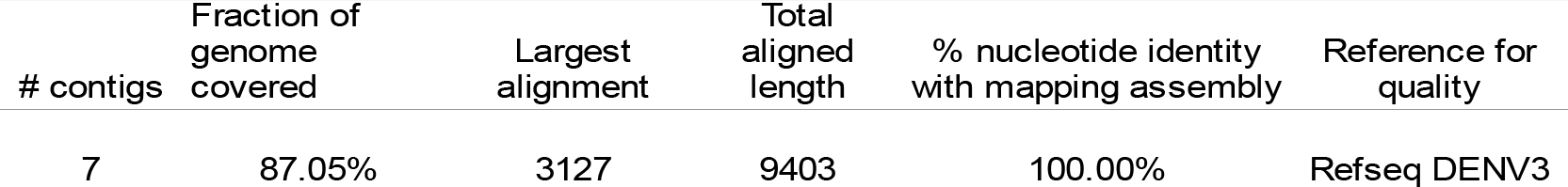
Assembly characteristics for de novo assembly. table shows the assembly characteristics by de novo assembly of sequences from sample F2 after quality assessment was performed using the QUAST tool. # contigs = No. of contigs.

| # contigs | Fraction of genome covered | Largest alignment | Total aligned length | % nucleotide identity with mapping assembly | Reference for quality |
| --- | --- | --- | --- | --- | --- |
| 7 | 87.05% | 3127 | 9403 | 100.00% | Refseq DENV3 |

The mapping-based assembly was used for phylogenetic analysis and submitted to GenBank as KX855927. Percentage nucleotide identity between this strain and the reference DENV3 genome (NC_001475.2) was 96.32% and with the closest DENV3 strain from India, 98.75%.

Phylogenetic analysis was carried out with BEAST2 using the coding sequence of KX855927 and 79 sequences selected as being similar to KX855927 using BLAST search against dengue genomes in the Virus Pathogen Database and Analysis Resource [34] (Additional File 3). The strain clusters with recent DENV3 sequences from India, China, and Singapore (Figure 2). This clade split from other DENV3 and other DENV3 genotype III strains around 15 years ago. The branch length of KX855927 is longer than most others in the tree, with an estimated divergence time of 13.86 years (with 95% Highest Posterior Densities between 12.94 and 14.83 years) from the closest Indian strain (Figure2). A maximum likelihood tree showed the same topology as the consensus tree from BEAST, although many clades had low bootstrap support (Additional File 4).

**Figure 2:**
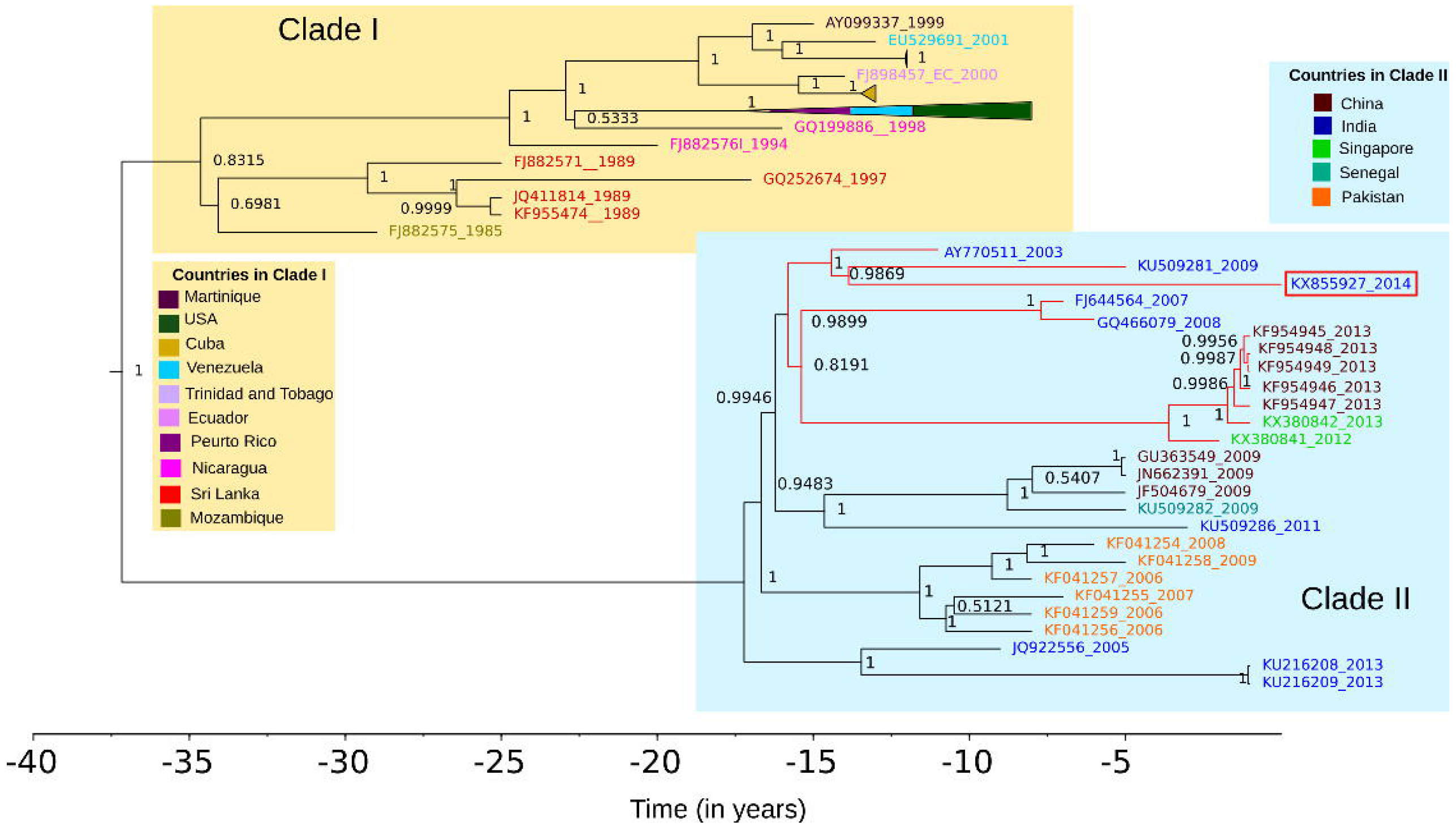
The sequenced strain KX855927 (2014) belongs to a recent Asian clade within genotype III. Figure shows the BEAST maximum clade credibility tree of top 79 BLAST matches to KX855927 The Indo - China - Singapore strain to which KX855927 (2014) is shown in red. All strains are represented by their GenBank IDs and coloured by country. For ease of visualization, a clade containing viruses from USA, Venezuela and Puerto Rico in Clade I has been collapsed (pyramids colored by country). The X axis represents time in years.

Both synonymous and non-synonymous substitutions were predicted throughout the genome as compared to the DENV3 reference sequence (Additional File 5). We aligned the envelope protein (E), of all the complete genomes from Indian strains against the parent strain used to derive the tetravalent dengue vaccine (CYD-TDV; Dengvaxia^®^, Sanofi Pasteur) (Figure 3). Multiple amino acid substitutions were predicted throughout the envelope protein and two additional stop codons (at amino acid positions 58 and 168) were observed in the DENV3 KX855927. Most of the amino acid substitutions were shared among all the Indian strains, while a D361E substitution was unique to the DENV3 strain reported here (Figure 3A). Nine out of 11 of the substitutions were mapped onto the surface of the envelope protein. Of these, six are in key antigenic sites, with three sites known to influence antibody binding (Figure 3B).

**Figure 3:**
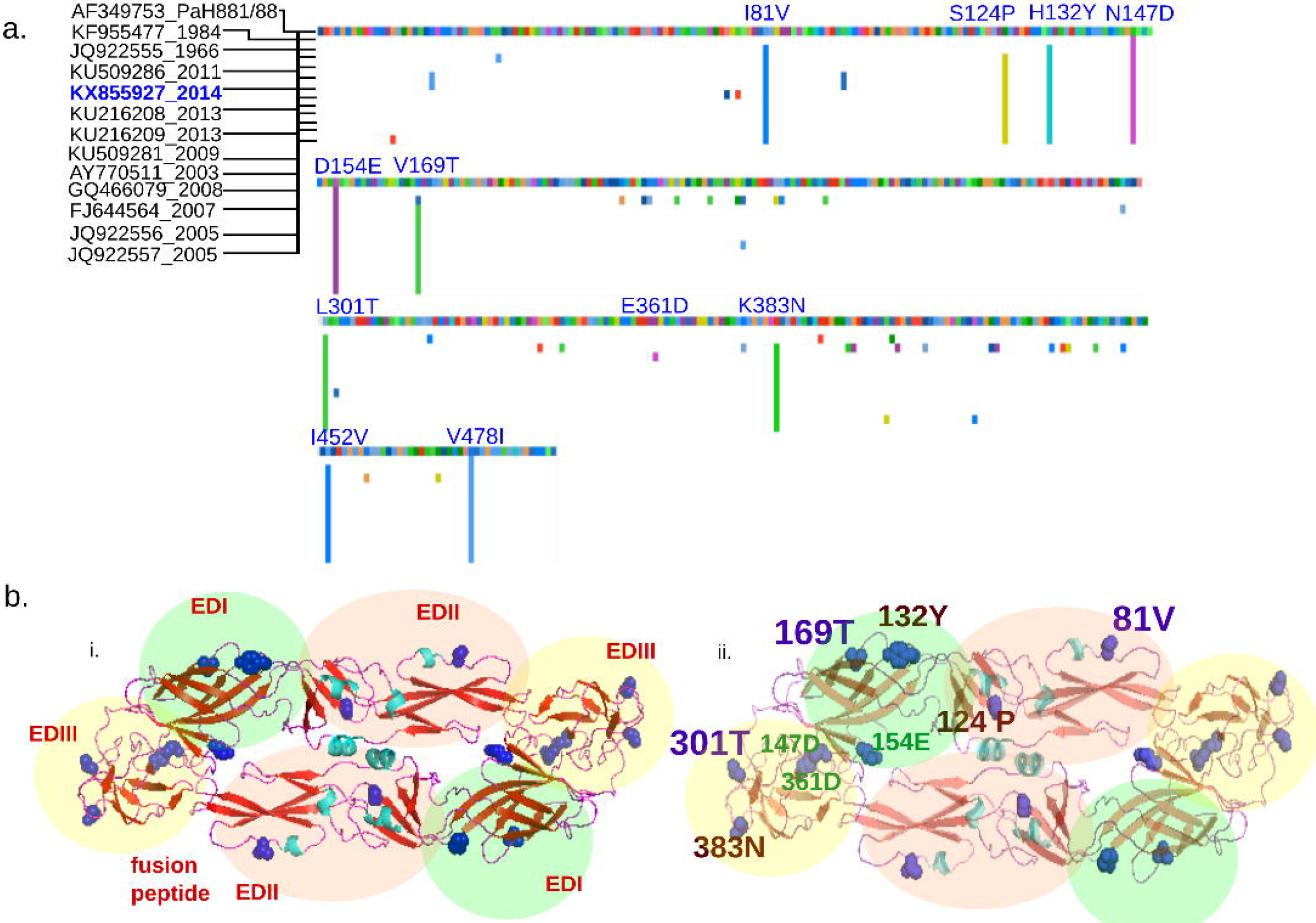
Shared amino acid substitutions in the envelope protein of Indian DENV3 strains differ from PaH881/8 a. Multiple sequence alignment of region coding for envelope (E) protein of dengue virus 3 from India were aligned to gi|13310784|gb|AF349753.1| DENV3 strainPaH881/88 polyprotein precursor, translated E genes. Numbers in the bracket represent year of sampling. Predicted amino acid changes compared to PaH881/88 are shown in colour. Position of substitutions present in the sequenced KX855927 strain are shown in blue. b. i) Cartoon structure of E protein KX855927 (2014)- dimer, homology modeled in SWISS-PROT with the domains shaded EDI (green), EDII (pink), EDIII (yellow),and labeled in red. ii) Cartoon structure of E protein KX855927 (2014)- dimer, homology modeled in SWISS-PROT showing the amino acid substitutions in KX855927 (2014) compared to the PaH881/8 in one of the dimers. In both cartoons, predicted substitutions are shown in blue (side-chains colored). Amino acid substitutions in violet are positions known to influence mouse monoclonal antibody binding. Positions in red are among 32 positions in the E protein predicted to be important for antigenicity.

The sequencing reads mapping to JEV from Sample F2 and F5 were assembled into contigs and used to check for potential alignment to other genomes in the NCBI nucleotide (NT) sequence database. The JEV sequences we identified, were specific to JEV (100% identity, 100% coverage of read) from a BLAST search (Additional File 6). The sequences were found to match non-structural protein 5 of JEV. Specific search against the dengue database for the contigs from the sample containing DENV3 sequences showed no similarity for contig 1 and some similarity to a dengue virus 2 sequence for contig 2 (83% identity, 97% coverage, Additional File 6).

The single DENV3 sequencing read found in sample F3 was identical to a sequencing read occurring with high frequency in sample F2. Therefore, we did not carry out any further analysis with this sequence read and suspect it to be a contamination.

## DISCUSSION

Here we sequenced a complete dengue genome from a clinical case of severe dengue fever, without the need to culture the virus, and in an unbiased manner. We believe that sequence based enrichment of viral sequences will enable recovery of complete genomes by lower depth sequencing from routine clinical samples [35].

We identified a low number of reads mapping specifically to Japanese encephalitis virus (JEV). JEV is known to cause fevers [5, 6, 36]. Further systematic analysis using a combination of polymerase chain reaction and IgM testing is required to ascertain how much JEV contributes to the acute fever burden in India. The low number of JEV reads obtained in both samples in which reads mapping to JEV were found, suggests there was not much active viral replication occurring. There are previous reports of detection of JEV sequences many months after infection [37]. The sequences we found could therefore be remnants of previous infection or may be the result of a infection from a mosquito bite which was checked by the immune system. The low number of reads in these cases mapped to the same gene (non-structural protein 5) (Additional File 6). This could reflect higher stability of some parts of the JEV RNA genome.

The results of metagenomic sequencing however do need to be interpreted with caution due to issues related to contamination [10, 11]. Contamination can occur in every step of the procedure, starting from sample collection, processing, sequencing and when multiple indexed samples are sequenced together, de-multiplexing (the process in which reads get assigned to a sample). This needs to be taken into consideration particularly when the number of sequences supporting the presence of a pathogen are low, when there is incomplete genome information, or the same sequence is present in all the samples including the controls. We have tried to mitigate this partially by use of controls - serum from a healthy individual collected at the same time and place, and a water sample processed in the same way as the clinical samples. However, we believe that independent methods are required to confirm novel/ unexpected findings by this method.

DENV3 has been shown to be re-emerging in India and has been responsible for severe outbreaks in different geographic regions, including in South America and Cuba [27, 38, 39]. The full-length DENV3 (KX855927) we describe here clusters into a clade containing DENV3 viruses from India and is related to an Indo - China - Singapore clade. We observed a longer branch length for this particular strain which could be the result of incomplete sampling of this clade or could indicate that this lineage is showing accelerated rates of molecular evolution [40]. This can be resolved in future studies by the addition of more sequence information, as more full-length dengue sequences from India become available in the databases.

While both synonymous and non-synonymous changes were observed throughout the DENV3 (KX855927) genome compared to the DENV3 reference sequence (NC_001475.2), the changes in the antigenic envelope protein are of particular interest. Neutralizing antibodies have been described against the envelope protein that target particular epitopes [26,41]. Critical amino acid residues that change antibody binding have also been described by others [26]. The results from our phylogenetic analyses are consistent with previous work tracing the emergence of new clade of DENV3 genotype III strains in India [39]. The ability of a dengue vaccine to elicit neutralizing antibodies against locally circulating DENV3 strains therefore needs to be evaluated in this light.

## CONCLUSION

Our work demonstrates the usefulness of a metagenomic approach to pathogen characterization starting with clinical sample. Currently this method is more expensive than routine molecular diagnostics, however we believe that the costs will decrease as the technology becomes more widely available, enabling its use in diagnostics. Our findings with dengue virus type 3 encourage the use of sequencing to track viral evolution and its relationship to the dengue disease landscape in India.

## MATERIALS AND METHODS

Description of samples - 5 patients (2 diagnosed as dengue fever (serum, F1-F2), 2 Rickettsial fever (Serum, F3, F5), 1 unknown fever (plasma, F4) presenting with febrile illness (Table 1: provides clinical characterization, treatment and outcomes) and 1 healthy control (serum, C1) sample were used in the study. The study was done after obtaining approval from the Institutional Ethics Committee of St. John's Medical College and Hospital(IEC Study Ref.No.5/2016). A waiver of consent was sought and obtained for the analysis as it was done on samples remaining after routine diagnostic testing and de-linked from identity of the patients.

Isolation of RNA - RNA was extracted using the Qiagen All-Prep kit, using 300-500 microliteres of serum/Plasma and lysed using 1 ml of lysis buffer. The remaining protocol was performed as recommended by the manufacturer. Eluted RNA was concentrated and used for sequencing reactions.

Sequencing - Sequencing libraries were prepared using the Ion Proton library preparation protocol. Indexing was performed using the IonXpress RNA Seq Barcode kit. Samples F1-4 and C1 were run on the same chip. F5 was run on a separate chip. Libraries were pooled to give equimolar concentrations of 10 pico molar. This was used in template preparation steps and RNA sequencing was performed using the Ion PI sequencing kit on the Ion Proton platform using the Ion PI^TM^ ChipV2 and Ion PI^TM^ Sequencing Kit V3.

Analysis of sequences - We aligned the sequencing reads to a database of all known viruses using the SNAP alignment tool [42]. All hits were verified using nucleotide BLAST sequence search and visualized using tools from the Dark Matter project (https://github.com/acorg/dark-matter.git). Reads aligning to human genome, human mRNA, rRNA large subunit and rRNA small subunit from the SILVA database were removed [43]. The aligned sequences were used as the input for assembly. De novo assembly was performed using the SPAdes (version 3.10.1) tool [44]. Quality assessment of the assembly was performed using the QUAST tool [45]. MIRA 4.0.2 was used for mapping based assembly, with the Genbank sequence NC_001475.2 for dengue virus 3 as the backbone for assembly and NC_001437 as the backbone for Japanese encephalitis virus [46]. Contigs were subjected to nucleotide BLAST using the online BLAST tool (https://blast.ncbi.nlm.nih.gov/Blast.cgi). The mapping based assembly of dengue 3 resulting from MIRA was manually checked for regions with low confidence using Gap5 [47]. Less that 30 nucleotides were found to have low confidence, of which 22 were in the 3'UTR end region. The files from the MIRA assembly together with the contributing reads are provided as Additional File 1. This sequence was submitted to GenBank as KX855927.

Phylogenetic Analysis - Phylogenetic analysis was performed with BLAST search hits to KX855927 in the VipR dengue virus database [34]. Only the coding sequence was used for the analyses. The alignment was visualized using the AliView software [48]. Nucleotide distances of KX855927 from other dengue viruses - reference sequence and the closest BLAST hit from India, were estimated using the MUSCLE alignment tool to create a percentage identity matrix [49]. The Generalized Time Reversible Model, namely GTR+I+G, GTR+I+G, GTR+G were found to be the best evolutionary models for codon positions 1, 2, and 3 respectively using PartitionFinder [50] where I represents invariant and G represents gamma, a shape parameter for the model. A previously estimated rate of substitution for dengue virus 3 =7.48 ×10^−4 subs/site/year (4.47E-4; 10.72E-4) was used to set a strict molecular clock [51]. The input XML file to BEAST (Version 2.4.6) [52] is provided in Additional File 3. Tracer was used to confirm sufficient sampling (ESS > 200 for all parameters). TreeAnnotator was used to generate the maximum clade credibility tree, where the node heights represent median height. Posterior probabilities for both the split of Clade I and II and the clade containing KX855927 were >95%. The tree was visualized using FigTree (http://tree.bio.ed.ac.uk/software/figtree/). The Maximum Likelihood tree was generated using thorough search and 1000 bootstraps using RaXML (Additional File 4) [53].

Analysis of envelope protein - Envelope (E) protein alignments for the DENV3 complete genomes from India were performed in AliView. Homology modeling was performed for the E protein of KX855927 using SWISS-MODEL and the best model was chosen for showing the substitutions. The protein surfaces as visualized by PyMOL (The PyMOL Molecular Graphics System, Version 1. 8 Schrödinger, LLC) are shown in light brown, amino acids found to be different in the KX855927 strain are colored by elements CHNOS.

The datasets supporting the conclusions of this article are included within the article and its additional files (1-6).

## DECLARATIONS

### Ethics, consent and permissions

The study was done after obtaining approval from the Institutional Ethics Committee of St. John's Medical College and Hospital(IEC Study Ref.No.5/2016). A waiver of consent was sought and obtained for the analysis as it was done on samples remaining after routine diagnostic testing and de-linked from identity of the patients.

### Consent for publication

Not applicable

### Availability of data and materials

The data sets supporting the conclusions of this article are included within the article and its additional files (1-6). The raw files from sequencing are not provided as these are metagenomic data sets with host information.

## Acknowledgements

Dr. Lisa Ng and Dr. Julian Hiscox for critical input and helpful discussions. Sreejayan Nambiar, Field application specialist, Thermo Scientific, for technical support during the sequencing.

## Author contributions

PC designed and performed the experiments, carried out the analysis and wrote the paper. MD designed the experiments, provided clinical material, carried out the experiments, contributed to interpretation of the data and wrote the manuscript. AS provided clinical material and designed the experiments. MG and SS designed the experiment and performed the library preparation and sequencing. DS, BM, TJ designed and performed the computational analysis. KT performed phylogenetic analysis and interpreted the data. TS interpreted the results and wrote the manuscript. SK designed the experiments and wrote the manuscript.

## Disclaimer

The funders had no role in the in any steps of experimental design, data collection or analysis or the decision to publish.

## ADDITIONAL FILES

Additional File 1- File format - ․maf (MIRA assembly format, can be converted to compatible file formats for viewing with Gap5, Consed and other genome editors)

Title of Data - dengue virus 3 assembly with contributing reads

Description - File contains consensus sequence and reads contributing to the assembly, it contains regions/ bases flagged by MIRA including 30 base positions which have low confidence.

Additional File 2 - File format - Document File (.doc)

Title of data - Percentage identity of DENV3 sequence assembled by different methods

Description - Table shows the percentage similarity between the de novo and mapping assemblies compared to NCBI reference sequences of dengue virus 1-4 and the closest Indian strain.

Additional File 3 - File format (.xml)

Title of Data: Template for BEAST

Description: The input file used for phylogenetic analysis using the BEAST program

Additional File 4 - File Format (.png)

Title of Data - Maximum Likelihood Tree from RaxML

Description of Data - Figure shows the maximum likelihood trees with bootstrap values, Genbank Ids and year of sequencing are shown on the tips. The data is coloured by country and some of the clades have been collapsed for ease of viewing.

Additional File 5 - File format - excel sheet (.xls)

Title of data - Single Nucleotide Polymorphisms in KX855927 with respect to NC_001475

Description - Description of the Single Nucleotide Polymorphisms in KX855927 with respect to NC_001475 as detected by the MIRA assembly program.

Additional File 6 - File format - excel sheet (.xls)

Title of data: BLAST results of Japanese encephalitis virus contigs from sample F2 and F5

Description- Contains the Top 5 nucleotide BLAST lists for the Japanese encephalitis virus contigs assembled from samples F2 and F5, against the nucleotide database, Flavivirus database and dengue virus database.

